# The glucocorticoid receptor ligand-binding domain confers drug-gated protein regulation in *C. elegans*

**DOI:** 10.1101/445833

**Authors:** Gabriela C. Monsalve, Keith R. Yamamoto, Jordan D. Ward

## Abstract

Controlling protein activity and localization is a key tool in modern biology. Mammalian steroid receptor ligand-binding domains (LBDs) fusions have been used in a range of organisms and cell-types to inactivate proteins of interest until the cognate steroid ligand is applied. Here, we demonstrate that the glucocorticoid receptor LBD confers ligand-gated control of a heterologous gene expression system (Q system) and the DAF-16 transcription factor in *C. elegans*. These experiments demonstrate provide a powerful tool for temporal control of protein activity, and will bolster existing tools used to modulate gene expression and protein activity in this animal.

## Introduction

The ability to temporally and spatially control gene expression and protein activity/localization are essential tools in modern genetics. Heterologous systems, such as Gal4/UAS have been widely used in *Drosophila melanogaster*, allowing researchers exquisite control over the time and tissue in which the transgene is expressed (Brand and Perrimon 1993). The Q system, which adapts the *Neurospora crassa* transcriptional activator, QF, and its repressor, QS, to gene activation via de-repression of QS by quinic acid, offers an additional system for transgene expression analysis, lineage tracing, and mosaic analysis in mammalian cells and in fly models (Potter *et al.* 2010; Riabinina *et al.* 2015). Other methods to control gene expression include tissue-specific expression of the Cre recombinase fused to the estrogen receptor ligand-binding domain (Feil *et al.* 1996), CRISPR interference and CRISPR activation (Qi *et al.* 2013; Gilbert *et al.* 2013; 2014), tetracycline-inducible gene regulation systems (TET ON/OFF)(Gossen and Bujard 1992; Gossen *et al.* 1995), and tags to stabilize or destabilize proteins, in some cases in an inducible manner (Dohmen *et al.* 1994; Banaszynski *et al.* 2006; Nishimura *et al.* 2009; Bonger *et al.* 2011; Holland *et al.* 2012; Morawska and Ulrich 2013; Zhang *et al.* 2015).

In *C. elegans*, the most common method to induce gene expression is to fuse heat shock promoters upstream of a gene of interest; following acute heat shock, gene expression is robustly induced (Stringham *et al.* 1992). Modifications to this approach include the FLP-OUT system (Voutev and Hubbard 2008), and the HSF-1 system, which allows for some tissue-specificity of gene induction (Bacaj and Shaham 2007).

These approaches have been powerful in *C. elegans*, but have some caveats: i) in addition to activating the transgene of interest, heat-shock also provokes regulation of heat-shock responsive genes and can affect cellular transcription and translation in ways that can cause unknown and undesired physiological effects; ii) tissue-restricted expression can be achieved, but requires specific genetic backgrounds; and iii) the system results in a pulse of target gene transcription rather than sustained transgene expression, although modifications such as the FLP-OUT may bypass this limitation. Bipartite gene expression systems, such as Gal4-UAS, have been historically lacking in *C. elegans*, likely because *C. elegans* is cultured at lower temperatures than S. *cerevisiae*, and thus the imported Gal4-UAS system proteins are less active at these lower temperatures. The Q system, which was imported from a fungus with a more similar growth temperature range to *C. elegans*, was adapted to *C. elegans* to control gene expression (Wei *et al.* 2012). Additionally, a temperature-robust Gal4-UAS system was also developed for *C. elegans* (cGal) by optimizing the three main components of the system to function between 15-25°C (Wang *et al.* 2016).

Previously, the *Neurospora crassa* QF/QUAS system was adapted to *C. elegans* to mark subclasses of neurons (Wei *et al.* 2012). In this system, the QF transcription factor binds to a response element (Q upstream activating sequence; QUAS) to drive transcription of a downstream transgene. Constitutive expression of a transgene can be repressed through co-expression of a repressor (QS), similar to how Gal80 is used in the Gal4-UAS system. Adding quinic acid relieves QS-mediated repression, and permits QF to drive expression of the transgene. Tissue-specificity can be obtained using split QF constructs and/or by expressing QF and QS using tissue-specific promoters, and the system functioned effectively using single-copy transgenes (Wei *et al.* 2012). Some limitations of the system are that QS and QF have to be expressed constitutively, and require a minimum of six hours for quinic acid to alleviate QS repression and allow transgene induction, with complete restoration of transgene expression taking 24 hours; this delay is not ideal for tightly controlling transgene induction in developmental contexts, or during dynamic and rapid cellular events.

Using the ligand-binding domains (LBD) from mammalian steroid receptors permits drug-inducible control of the Cre and Flp recombinases and the Gal4-UAS system (Picard *et al.* 1988; Logie and Stewart 1995; Metzger *et al.* 1995; Kozlova and Thummel 2002). We were interested in using steroid receptor LBDs would allow for both robust repression of fused factors and rapid alleviation of this repression upon ligand addition. We therefore wished to explore whether the human glucocorticoid receptor LBD could be used to regulate two proteins in *C. elegans:* the QF transcriptional activator and the DAF-16/FOXO transcription factor.

## Materials and Methods

### Strains and Maintenance

All *C. elegans* strains were cultivated on NGM plates using standard methods. Animals were maintained at 20°C, unless otherwise denoted. The following mutant and transgenic strains were used in this study: **N2** (Wild-type, Bristol)(Brenner 1974); **EG5003** *(unc-119(ed3)* III; *cxTi10882* IV); **CB6193** *(bus-8(e2885)* X); **PS5980** *(him-5(e1490) syIs197[hsp16-41::LIN-3C* cDNA + myo-2p::dsRed + *pha-1(+)*] V; outcrossed 4X); **CF4087** *(daf-2(e1370)* outcrossed 12X). Strains generated for this study are listed in Supporting Information, File S1, Table S1.

### Cloning of dex-inducible constructs

Plasmids containing the full-length coding sequences for the QF activator, the 4X-repeat of the *quas* response element, and the QS repressor (pXW83, pXW82, and pXW09 respectively), and their associated selection markers, were described previously (Wei *et al.* 2012). The nucleotides 2967-3737 from cDNAs encoding for the human glucocorticoid receptor alpha (GR), which correspond to its ligand-binding domain, was amplified from pFastBacGRα-6XHis (Yamamoto Lab, UCSF), and fused in frame to the C-terminal end of full-length coding sequence for QF using In-Fusion ligation (Clontech; Catalog #638909). Gateway cassettes were engineered upstream of QF-GR coding region (pGM32DEST) or QS coding region (pGM48DEST), or downstream of the QUAS response element and minimal delta pes-10 promoter (pGM34DEST) using In-Fusion ligation. Using directional primers, approximately 0.5-2kb upstream of the start codon of *atf-8, eef-1A.1, egl-17, and pro-1* genes was PCR-amplified from N2 genomic DNA.

Similarly, the cDNAs from *peel-1* were PCR amplified from the pMA122 plasmid (Addgene, plasmid #34873), and the *lin-3c* gene was cloned from the pBlueScript_lin-3c plasmid (a generous gift from Cheryl Van Buskirk, Cal State Northridge). Blunt PCR fragments were then TOPO-cloned into the pENTR/D-TOPO Gateway entry vector (Invitrogen; Catalog #K240020), and the resultant clones were recombined with the appropriate destination vectors. All recombinant plasmids were subsequently injected into *unc-119* mutant animals, with a combination of 40 ng/μl of each recombinant plasmid, and pBlueScript, for a total of 100 ng/μl DNA, with the exception of *quas::peel-1*, which was injected at 15 ng/μl. Transgenic progeny that were both phenotypically wild-type and express fluorescent mCherry were subsequently isolated and tested for dex-induced gene expression. In-Fusion ligation (Clontech) was utilized to fuse the GR LBD from pGM32DEST in frame and upstream of both eGFP and DAF-16A.

### Preparation of Ligand Stocks

100mM (1000X) dex (Sigma; Catalog #D1756) stocks were prepared by solubilization in DMSO, followed by 0.2um filter sterilization; stocks were stored at −80°C until use. 1000X dex was then added to liquid NGM (cooled to 55°C) along with salts and cholesterol to a final concentration of 100μM dex. Alternatively, dex was spread on top of existing NGM plates if the volume was known, or added to a worm/M9 slurry to a final concentration of 100μM. Stock solutions of 300mg/mL of quinic acid (pH 6-7) were similarly added to plates or worm/M9 solutions, as previously described (Wei *et al.* 2012). Freshly prepared 10mM fluorescein-dexamethasone stocks (ThermoFisher; Catalog #D1383) were solubilized in ethanol, sterilized through a 0.2um filter, and stored at 4°C in the dark until use.

### Molecular Biology

For the extraction of total RNA, selected animals were placed in Trizol (Invitrogen; Catalog #15596026) and subsequently lysed by rapid freeze cracking. The RNeasy extraction kit (Qiagen; Catalog #74104) was used to remove contaminating genomic DNA and isolate total RNA. The iScript cDNA synthesis Kit (Bio-Rad; Catalog #170-8891) was then used to synthesize cDNAs. To measure the levels of transcripts, master mixes using cDNA templates, SsoAdvanced Universal SYBR Green Mix (Bio-Rad, Catalog #1725270), and gene-specific primers were used. All reactions were performed on the Bio-Rad CFX Connect Real-Time PCR Detection System instrument (Bio-Rad; Catalog #1855200). The primers used in this study are listed in Supporting Information, File S1, Table S2.

### Fluorescence Microscopy

All images were prepared for publication using Image J, Adobe Photoshop and Adobe Illustrator. To score GFP expression, we transferred 10-20 mixed-staged animals to NGM plates treated with vehicle or 100μM dex. Populations were cultivated at 25°C, and fluorescence was assessed repeatedly every hour at 5X to 10X magnification using the FSM25 Kramer Fluorescence Microscope, connected to an X-Cite 120Q fluorescent light source. GFP and mCherry fluorescence were performed blind to the experimental conditions, and at least three independent experiments were performed.

The remaining microscopy was performed on a Zeiss Axioplan 2 fluorescent microscope attached to Xenon excitation lamps and green/red fluorescence filter sets. Images were captured with the attached Hamamatsu Orca ER camera controlled by Micro-Manager, an open-source microscopy software. For these experiments, animals were paralyzed with 10mM levamisole, and mounted onto a 2% agar pads. For the fluorescein-dexamethasone (F-dexa) experiments, mixed-staged animals were washed three times for 15 minutes with M9 solution, soaked in an M9+0.05% gelatin solution containing 1-100μM F-dexa, and rotated on the benchtop for two hours. Live populations were then washed an additional three times with the M9 solution before imaging. In populations expressing the *hsp-16.48* promoter, mixed-staged animals were gently washed off of NGM plates, and resuspended in M9 with vehicle or 100μM dex. A 30min, 33°C heat shock was performed in a thermocycler, and animals were incubated at room temperature for an additional 3.5 hours on a bench top rotator before imaging. To assess vulva morphology, populations were synchronized by extracting eggs from gravid adults by alkaline lysis followed by hatching M9+0.05% gelatin for 24-48 hours at 25°C. Next, L1 hatchlings were released from the starvation-induced L1-diapause by feeding, and subsequently grown to the mid-L4 larval stage. L4 larvae were then washed off of plates, and treated with vehicle or 100μM dex for three hours rotating on a bench top before imaging. Vulval morphology was assessed in mCherry positive animals in late L4 larvae or young adults, at 16X-25X magnification.

### Behavioral Assays

For measurements of behavioral adult quiescence, active young adults were transferred into either vehicle or 100μM dex solutions in M9 buffer in a total volume of 100μl, and then incubated on a rotator for two hours at room temperature. Animals were transferred to an NGM plate seeded with OP50 *E. coli* for 30 minutes at room temperature, and then individual adults were assessed for behavior. Behavioral quiescence was defined by two criteria: 1) the absence of pharyngeal pumping for 60 secs, and; 2) the cessation of body movements for at least 30 secs, as previously described (Van Buskirk and Sternberg 2007; Monsalve *et al.* 2011; Hill *et al.* 2014; Nelson *et al.* 2014). Behavioral assessments were done with at least three independent trials on separate days, in parallel to the *hsp::lin-3c* positive control strain (PS5970), and blind to the experimental conditions.

### Statistical Analysis

Relative gene expression was calculated by determining the fold-change variation over control (vehicle) samples using the Comparative C_T_ method (Schmittgen and Livak 2008). Regression analysis was utilized to determine C_T_ values, and the mean C_T_ value from reactions preformed in triplicate was used to determine the average fold-change from the *ama-1* internal control. Error bars were calculated using the error propagation of standard deviations to the logarithmic scale. The comparisons between vehicle and dex-treated animals (and thus limited to two conditions) were performed using an unpaired, two-tailed Student’s T-test, or a Chi-square test. All p-values were calculated using the GraphPad Prism 6.0 software.

### Data Availability

Strains generated in this study will be available via the Caenorhabditis Genetics Center (CGC) at the University of Minnesota-Twin Cities and/or through direct request to J.D.W. The destination vectors described in this study (pGM32DEST, pGM34DEST, and pGM48DEST) will be available via Addgene.

## Results

### A drug-inducible expression tool for *C. elegans*

To engineer a heterologous, drug-inducible gene expression system for *C. elegans*, we modified the QF transcriptional activator by fusing the ligand-binding domain (LBD) from the human glucocorticoid receptor (GRα; NCBI Gene ID: 2908) at the C-terminus (QF- GR; Figure 1A). We chose the glucocorticoid receptor as *C. elegans* does not have any clear orthologs (Antebi 2015), and ligand binding by the GR LBD is very specific, whereas estrogen receptor ligand-binding domain exhibits promiscuity (Eick *et al.* 2012). QF-GR should be inactive until the synthetic, non-metabolized GR ligand dexamethasone (dex) is added (Scherrer *et al.* 1993). QF-GR, was cloned into a plasmid with a contiguous splice leader (SL) and an mCherry reporter, which marks cells in which QF-GR is expressed. Though designed for Gateway cloning (Figure S1A), our system can be easily converted for any ligation-dependent or -independent cloning pipeline. Transgenes are inserted downstream of a regulatory element containing a delta *pes-10* minimal promoter and a 4X repeat of the QF response element (QUAS) (Seydoux and Fire 1994; Wei *et al.* 2012).

**Figure 1:**
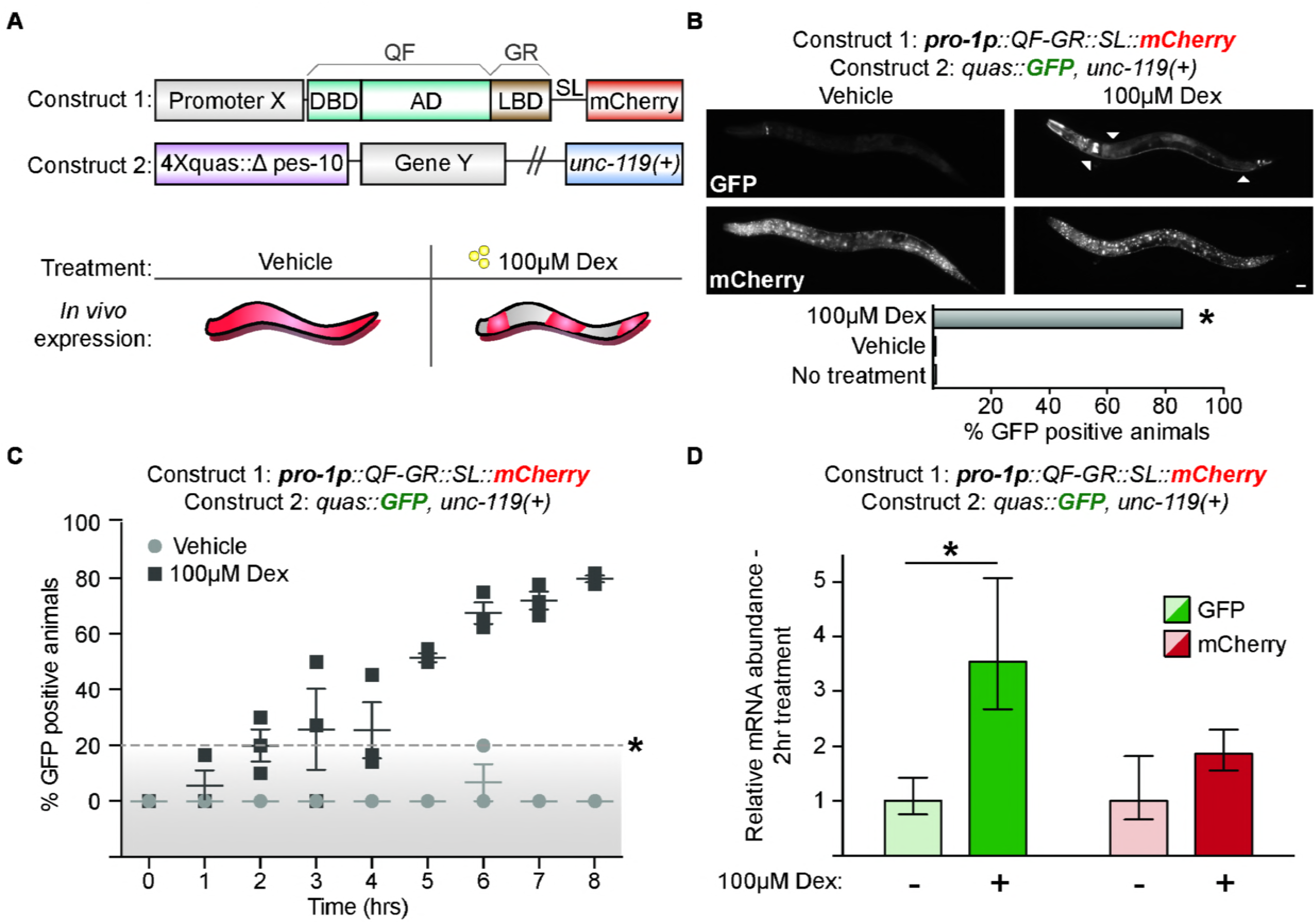
A drug-inducible gene expression system for *C. elegans*. **(A)** Schematic of a drug-inducible system for *C. elegans*, which consists of QF-GR transcriptional activator and the QUAS reporter plasmids. QF-GR consists of the full-length QF DNA-binding domain (DBD) and Activation domains (AD), fused to the ligand-binding domain (LBD) from the human glucocorticoid receptor α (GRα). The splice leader (SL) links mCherry expression to the activity of Promoter X, and marks for the expression of the array. The QUAS reporter consists of four tandem repeats of the QUAS response element upstream of the minimal delta *pes-10* promoter and Gene Y. **(B)** GFP and mCherry expression in animals containing QF-GR under the control of the ubiquitous *pro-1* promoter and a QUAS::GFP reporter. Fluorescent micrographs depict representative animals with the indicated transgenes cultivated on either vehicle or 100μM dex plates for 24 hours. Arrowheads denote simultaneous expression of GFP in neuronal and intestinal tissues upon dex-treatment. Ubiquitous mCherry expression is observed in a dex-independent manner. Scale bar denotes 20 microns. The percentage of GFP-positive animals observed is denoted (N ≥ 54 animals for each condition; *p<0.001, Chi-square Test). **(C)** A time course scoring GFP expression in animals carrying the indicated transgenes. Mixed-staged animals were cultivated on vehicle or 100μM dex plates, and GFP expression was scored hourly. Each timepoint denotes the mean percentage of GFP-positive animals (±SEM) from at least three independent experiments with more than 21 animals for each timepoint. Points above the dashed line denote a significant difference from vehicle-treated animals (*p<0.01, T-Test). **(D)** Average fold change of expression of dex-treated vs vehicle-treated populations with the indicated transgenes. qRT-PCR measurement of GFP and mCherry transcript level after two hours of vehicle (-) or 100μM dex-treatment (+) (±SEM from three biological replicates performed in triplicate; *p<0.05, T-test).

### Glucocorticoids are absorbed by *C. elegans*

We first tested if the GR ligand, dex, could be effectively absorbed into live animals. We used a fluorescein-dex (F-dexa) conjugate previously used to monitor the uptake of dex in other systems (Maier *et al.* 2005). As uptake of molecules in *C. elegans* can occur via oral ingestion and/or cuticle penetration (Valdes *et al.* 2012; Lee *et al.* 2015), we assessed uptake of F-dexa in wild-type (N2) animals, and in *bus-8* mutants which have a compromised cuticular barrier (Partridge *et al.* 2008). Mixed-stage populations were cultured with increasing amounts of F-dexa for two hours and localization of F-dexa was visualized using fluorescence microscopy. In both WT animals and *bus-8* mutants, we observed F-dexa exclusively in the pharyngeal and intestinal lumen in 97-100% (n=1644) animals; notably, fluorescence was not detected in the epithelia of treated animals (Figure S1B), and no fluorescence was observed in vehicle-treated populations (n=419, *p<0.00001, Chi-square test). With increasing concentrations of F-dexa, fluorescence was detected strongly and predominately within the intestinal lumen. F- dexa labeling of intestinal tissues was only observed at the highest tested concentration (100μM). These results indicate that 100μM of F-dexa is sufficient for its uptake into *C. elegans* intestinal cells, and further suggest that the intestine may be the tissue through which dex is absorbed.

### The GR LBD makes QF ligand-gated

We next asked whether dex and the GR LBD could gate the activity of the heterologous QF-GR protein. We expressed QF-GR ubiquitously using the strong *pro-1* promoter (Hunt-Newbury *et al.* 2007), and a GFP reporter was included under control of the 4xQUAS element; a cistronic mCherry cassette was used to mark cells in which the promoter was expressed. We observed induction of GFP 24 hours after cultivation with 100μM dex, but not after vehicle-treatment (Figure 1B). GFP expression was detected in 86% (n=125) of animals expressing the mCherry reporter, as compared to 0.9% (n=115) of vehicle-treated animals, and 0% (n=54) of animals with no drug treatment (*p<0.001, Chi-square test). GFP was detected robustly and predominately in neuronal, pharyngeal, and intestinal tissues (Figure 1B), and we also observed additional, modest GFP expression in the hypodermis and muscles of other animals (data not shown); however, we did not observe GFP nor mCherry expression in gonadal tissues. We did notice muted, yet detectable expression of GFP in a pair of unknown head neurons in both vehicle and dex-treated populations; this background pattern of expression was consistent to that observed in animals expressing only the reporter gene (Figure 1B and data not shown). The spatial pattern of GFP and mCherry expression was similar across larval development and in adults. However, we noted one distinct difference in embryos: while mCherry expression was detected in eggs, GFP expression was never observed in vehicle or dex conditions (data not shown). Finally, to determine if sustained cultivation with dex was toxic to animals, we measured the brood sizes of N2 and transgenic animals cultured on NGM plates supplemented with vehicle or 100μM dex. We observed no significant differences in the brood sizes between vehicle and dex-cultured populations: vehicle-treated N2 animals had an average brood size of 225±7, as compared to 222±13 of dex-cultured wild-type animals (mean number of progeny per animal ± SEM; n=10 in each condition; p=0.85, T-test). We similarly detected no significant differences in the brood sizes of pro-1p::QF-GR transgenic animals: vehicle-treated animals had an average brood size of 116±8, as compared to 98±13 of dex-cultured transgenics (mean number of progeny per animal ± SEM; n=10 in each condition; p=0.29, T-test). The *unc-119(ed3)* background mutation of the *pro-* 1p::QF-GR, *quas::GFP; unc-119(+)* transgenics likely contributed to approximate twofold decrease in brood sizes observed between the rescued transgenics and wild-type animals (*p<0.05, ANOVA test). Finally, we did not observe any abnormalities in morphology, foraging behavior, or developmental timing of the animals cultured in dex (data not shown).

Next, we performed a time course to assess how quickly GFP was detectable by fluorescence microscopy following dex exposure. Mixed-stage animals were cultivated on plates containing either vehicle or 100μM dex, and scored hourly for GFP and mCherry expression. After two hours of drug-treatment, 20% (n=25) of animals exhibited GFP fluorescence, as compared to no vehicle-treated animals (n=21; *p<0.03, T-Test) (Figure 1C). Using qPCR to measure GFP and mCherry transcript levels, we observed a 3.5-fold increase in GFP levels in dex-treated animals, as compared to vehicle-control animals (Figure 1D); mCherry mRNA levels did not change significantly within the same dex and vehicle-treated populations. In contrast, after 30 minutes of ligand exposure, we did not observe any significant changes in GFP transcript levels (Figure S1C). GFP was detected initially in the nervous system, and most predominately in unidentified tail neurons and the ventral cord. Notably, both the percentage of GFP-positive animals, and the tissue-distribution of GFP expression increased over time, with 80% of animals (n=38) expressing GFP in multiple tissues after eight hours of cultivation on dex-treated plates, as compared to 0% (n=27) of vehicle-treated populations (*p<0.001, T-Test). We similarly expressed QF-GR from an alternative ubiquitous driver, *eef-1A.1*, and observed dex-specific GFP induction in transgenic animals (Figure S1C). QF-GR expression from the *eef-1A.1* promoter drove significant reporter expression predominately in hypodermal, intestinal, and vulval tissues after two hours of dex-exposure in 27% (n=26) of animals, as opposed to 0% (n=22) of vehicle-treated animals (*p<0.04, T-Test) (Figure S1D). In contrast to transgenics expressing QF-GR from the *pro-1* promoter, we did not observe GFP in the nervous system in eef-1A.1::QF-GR animals. Moreover, expression of GFP in the intestinal, hypodermal, and vulva tissues appeared synchronously, and the intensity of GFP fluorescence increased with longer exposures of dex (data not shown). Together, these results indicate QF-GR permits ligand-gated transgene expression in most major *C. elegans* tissues within two hours of dex exposure, and that promoter choice is an important experimental consideration.

### QF-GR allows tissue-specific transgene expression

Having established that the GR LBD inactivated QF and that dex exposure allowed for QF activation, we next asked if QF-GR could drive tissue-specific transgene expression. We expressed QF-GR in vulval and hypodermal tissues, using the tissue-specific *egl-17* and *atf-8* promoters, respectively (Hunt-Newbury *et al.* 2007; Ward *et al.* 2013). Transgenic animals were cultured in vehicle or 100μM dex for three hours, and ratio of GFP-positive animals to mCherry-expressing animals was scored. We observed that 41% (n=26) of dex-treated *egl-17p::QF-GR* transgenics expressed GFP in the vulva, as compared to 0% (n=25) of vehicle-treated animals (Figure 2A). Similarly, hypodermal expression of QF-GR from the *atf-8* promoter resulted in 56% (n=41) of animals expressing GFP with 100μM dex, as compared to 22% of vehicle-treated animals (n=44) (Figure 2B). In both the atf-8p::QF-GR and *eef-1A.1p::QF-GR* transgenics, we observed a higher proportion of vehicle-treated animals expressing GFP after three hours of treatment compared to the pro-1p::QF-GR and *egl-17p::QF-GR* animals (Figure 2B and Figure S1D). These results demonstrate that dex can regulate the tissue-specific activity of QF-GR and its downstream reporter expression. Moreover, these experiments reveal that particular tissue-expression of QF-GR may affect its basal activity, as previously reported for ligand-activated GAL4 in *Drosophila* (Osterwalder *et al.* 2001; Duffy 2002).

**Figure 2:**
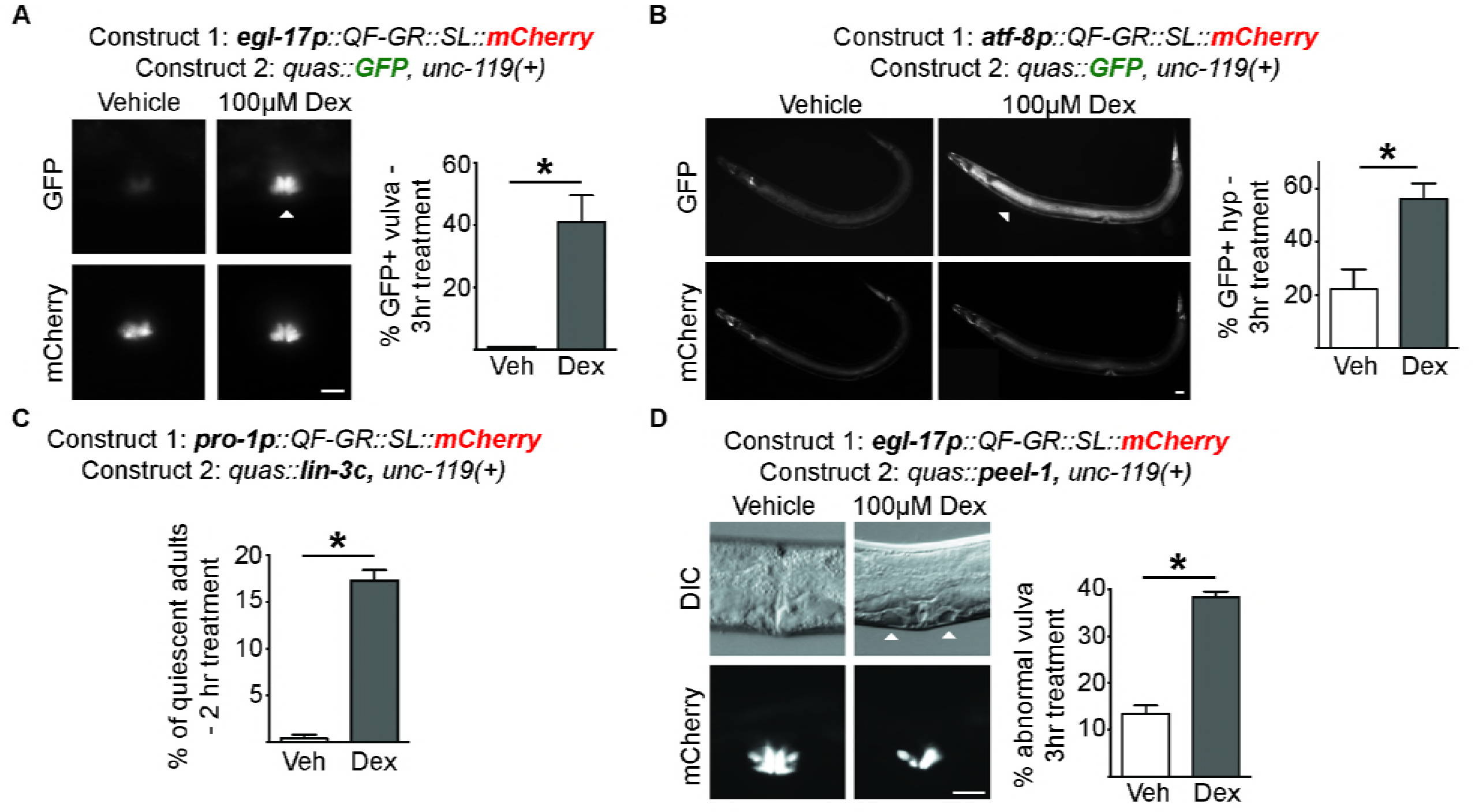
The QF-GR activator drives expression of target genes in various tissues. **(A-B)** Tissue-specific expression from the QF-GR activator in the **(A)** vulva or **(B)** hypodermis using the *egl-17* or *atf-8* promoters, respectively. The QUAS reporter contains the *GFP* gene. Representative fluorescent micrographs of GFP and mCherry expression from animals carrying the indicated transgenes, and treated with vehicle or 100μM dex for three hours. Arrowheads denote the tissue-specific expression of GFP; scale bar represents 20 microns. *(Right)* To the right of each set of images is a bar graph denoting the average percentage of GFP-positive animals after an acute three hour vehicle or dex treatment (±SEM; n ≥ 25 animals; *p<0.02, T-Test). **(C)** Induction of *lin3c/EGF* induces behavioral quiescence in adult animals. QF-GR::SL::mCherry was expressed using the ubiquitous *pro-1* promoter and the QUAS reporter contains the *lin-3c* gene. The graph depicts the percentage of quiescent adult nematodes after two hours of dex treatment. Behavioral quiescence of each animal was scored by two criteria: 1) the cessation of body movements for at least 30 secs, and 2) the absence of pharyngeal pumping for 60 secs (n ≥ 180 adults; *p<0.005, T-Test). **(D)** QF- GR::SL::mCherry was expressed using the *egl-17* promoter, which is expressed primarily in vulval cells, and the QUAS reporter contains the *peel-1* toxin gene. *(Left)* Induction of the *peel-1* gene by dex in vulval cells. Representative Nomarski micrographs of vulvae in late L4 larvae treated with vehicle or 100μM dex for three hours. Arrowheads mark holes in the vulva of a dex-treated animal due to presumed cell death; fluorescent images depict the vulva-specific expression of mCherry. Scale bar denotes 20 microns. *(Right)* The graph depicts the average percentage of L4s and young adults with abnormal vulval morphology after an acute three hour vehicle or dex treatment (±SEM; n ≥ 279 animals for each condition; *p< 0.0001, T-Test).

We next wished to move beyond fluorescent reporters and test whether QF-GR could be used to express transgenes to modify cell and animal physiology. Behavioral quiescence in *C. elegans* is characterized by cessation of feeding and movement, and delayed responses to noxious odors and touch (Trojanowski *et al.* 2015). Lethargy in animals can be triggered, in part, by the EGF ligand, encoded by the *lin-3c* gene (Van Buskirk and Sternberg 2007; Hill *et al.* 2014). We placed *lin-3c* cDNA under the control of the QUAS element and expressed QF-GR ubiquitously using the *pro-1* promoter (Figure 2C). We scored transgenic animals exposed for two hours to either vehicle or 100μM dex, and then assessed the behavioral quiescence of adult animals as described in Methods. We observed an approximate 30-fold difference in behavioral quiescence of dex-treated adults, as compared to vehicle-exposed adults (*p<0.005, T-Test). Specifically, 17% (n=181) of adults exhibited behaviors consistent with *lin-3c-* induced quiescence after a two hour treatment of dex, as compared to 0.4% (n=180) of vehicle-treated animals (Figure 2C). These observations indicate that the forced expression of *lin-3/EGF* can be achieved with dex-inducible QF-GR.

A powerful application of Gal4-UAS in *Drosophila* is for the conditional cell ablation by expressing death genes, such as Reaper (White *et al.* 1994). In *C. elegans*, the *peel-1* gene encodes a sperm-specific toxin, which destroys both germ and somatic cells in a cell autonomous manner. Accordingly, ubiquitous overexpression of the *peel-1* gene results animal lethality (Seidel *et al.* 2011). To test if our system could be used to conditionally induce animal lethality, we attempted to induce multi-tissue expression of *peel-1* using pro-1p::QF-GR; however, we were unable to propagate the transgenic lines due to toxicity of the arrays, even after multiple trials with various gene dosages (data not shown). Therefore, we then asked if, alternatively, our system could be utilized to drive cell death in a tissue-specific fashion, as was done in other organisms (Gerety *et al.* 2013; Obata *et al.* 2015) To test this, we expressed QF-GR using the *egl-17* promoter, which drives expression of QF-GR primarily in vulval – and some hypodermal – tissues. Synchronized mid L4 stage animals were treated for three hours with 100μM dex or vehicle and scored morphologically at the late L4 and young adult stages. Abnormal vulvas were observed in 39% (n=282) of dex-treated L4s, as compared for 14% (n=279) of vehicle-treated L4s (Figure 2D). Continued treatment with dex resulted in lower brood sizes (data not shown), suggesting that damage to the vulva from the *peel-1* induction affected the overall fecundity of the population. Taken together, these results demonstrate that our system permits drug-inducible transgene expression of complex behavioral and morphological processes, and underscores that promoter choice affects basal expression of QF-GR.

### The QS repressor and quinic acid together restrain the activity of QF-GR

*Neurospora* QS protein inhibition of QF activity is relieved by addition of quinic acid (Wei *et al.* 2012). To test whether QS could inhibit the activity of QF-GR, we co-expressed QF-GR and QS, and monitored QUAS::GFP reporter activity. As expected, we failed to detect GFP fluorescence in animals cultured in either vehicle only or vehicle plus quinic acid (n≥57) (Figure 3B). Following dex-treatment in the absence of quinic acid, we observed GFP expression in only 0.9% and 1.0% (n≥85) of animals. However, when transgenic animals were co-treated with both dex and quinic acid, we observed that 40% and 54% (n≥80) of animals expressed GFP (Figure 3B). Using qRT-PCR to assess induction of transgene transcripts, after a two hour vehicle-treatment, the QF-GR activator increased GFP levels approximately 28-fold above basal expression; addition of 100μM dex increased GFP expression to 98-fold above basal expression (Figure 3). As before, QS suppressed the expression of GFP mRNAs, as transcript levels were only 1.7-fold higher in both vehicle and dex-treated populations relative to basal expression (Figure 3C). After pre-treatment with quinic acid for approximately 24 hours, we measured a small, but significant 2.3-fold increase in GFP transcript expression after an acute, two hour dex-treatment. This dex-specific amplification was further intensified after a 49 hour treatment with both dex and quinic acid. We identified an 11-fold increase in GFP mRNAs in dex-treated animals, and only a 1.2-fold increase in GFP transcripts in vehicle-treated animals, as compared to the basal activity from the *quas::GFP* reporter. These data suggest that QS can repress transcriptional activity even in the presence of dex, and that QS de-repression and dex activation provide dual regulation, analogous to numerous bacterial operons that are under both negative and positive control.

**Figure 3:**
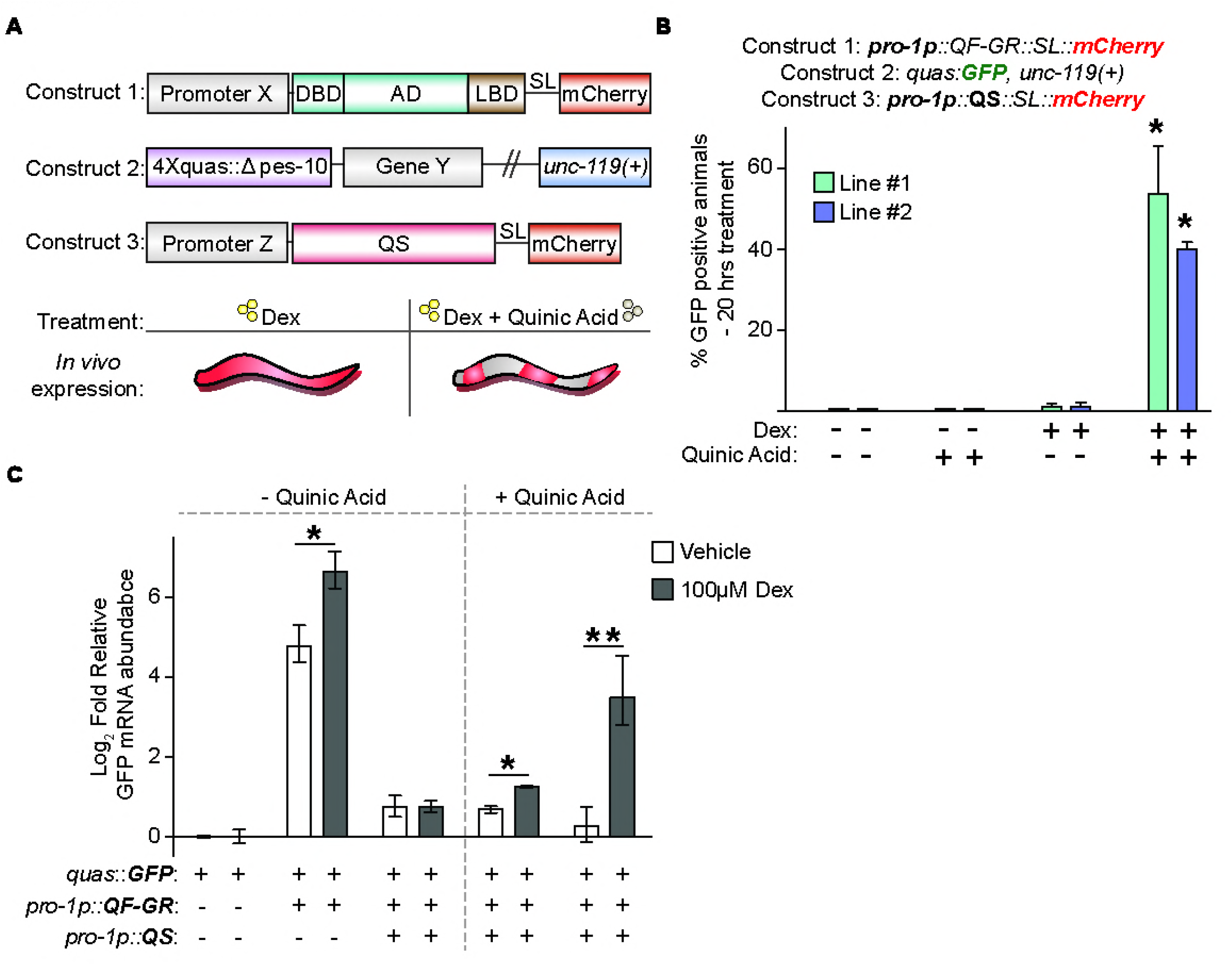
The QS repressor blocks activity of QF-GR. **(A)** Construct and experimental schematic of the addition of QS to the dex-inducible gene expression system. The composition of constructs 1 and 2 are described in Figure 1. Construct 3 encodes for the QS repressor, and a splice leader that links the mCherry open reading frame to mark expression of the array. The addition of the QS repressor restricts the activity of the QF-GR activator. De-repression of QS is achieved by the addition of quinic acid; therefore, only in the presence of both dex and quinic acid will QF-GR drive target gene expression. **(B)** The mean percentage of GFP-positive animals in animals carrying the indicated transgenes. QF-GR::SL::mCherry is expressed using a ubiquitously expressed *pro-1* promoter and QUAS drives a GFP reporter. *pro-1p* also drives the expression of the QS repressor, which restricts the activity of QF (Wei *et al.* 2012); de-repression achieved by the addition of quinic acid. The splice leader links mCherry expression to QS expression to mark sites in which the transgene is expressed. Error bars represent the SEM from two independent experiments (n ≥ 57 animals for each condition; *p<0.05, T-Test, as compared to both vehicle-treated (-) populations). **(C)** qRT-PCR measurement of relative GFP transcript levels in animals carrying the indicated transgenes. The graph presents mean, log2 fold change of expression of the indicated transgene/drug combinations relative to basal expression of vehicle-treated animals carrying the *quas::GFP* reporter (no QF-GR or QS transgenes). Animals exposed to 7.5mg/ml quinic acid were pretreated for 24 hours prior to dex treatment. Error bars indicate the SEM from three biological replicates performed in triplicate. The single asterisk (*) reflects the comparison by T-Test of animals treated with quinic acid for 24 hours and then vehicle or dex for two hours; double asterisks (**) represent the comparison by T-Test of animals treated with quinic acid for 24 hours and then vehicle or dex for an additional 25 hours (for both * and **, p < 0.05).

### The GR-LBD adduct modulates DAF-16 and GFP activity *in vivo*

Finally, we wished to determine whether the GR LBD could be more broadly used to control the activity and localization of other *C. elegans* proteins. We fused the GR LBD adduct to eGFP under the control of a heat-shock-inducible promoter *(hsp-16.48)*, which drives expression predominately in the muscle and hypodermis (Stringham *et al.* 1992). After a four hour treatment with 100μM dex, GFP expression was detected only in the hypodermal nuclei of 100% (n=78) of animals; in contrast, GFP was visible in both the cytoplasm and nucleus in 55% (n=94) of vehicle-treated animals (Figure 4A). This result suggests that the GR LBD adduct may function to restrict localization of the linked protein to the cytoplasm in the absence of ligand, and upon dex-binding, translocation to the nucleus occurs more readily (Picard and Yamamoto 1987).

**Figure 4:**
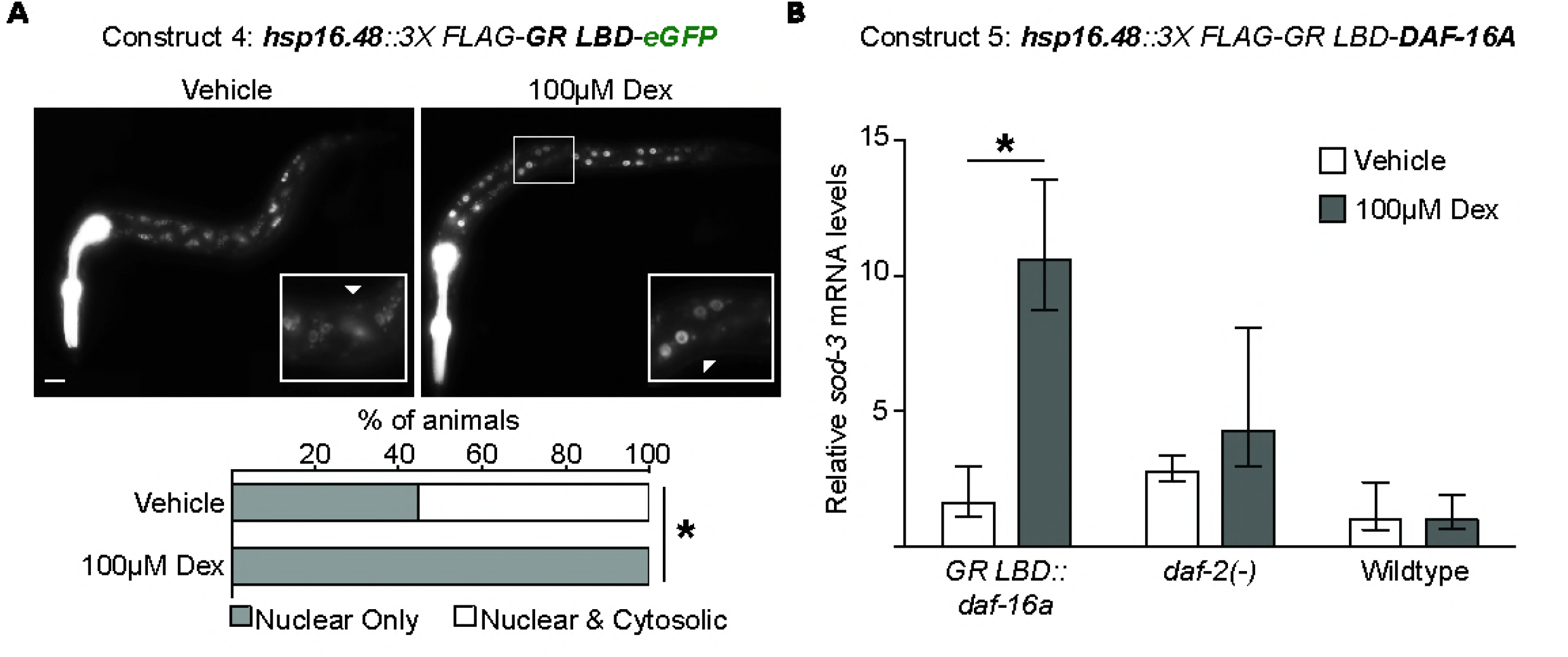
The GR-LBD adduct modulates protein activity *in vivo*. **(A)** Representative fluorescent micrographs of L1 transgenic animals expressing the GR LBD fused in frame to the N-terminus of GFP, and under control of the heat-shock inducible *hsp-16.48* promoter. Animals were heat-shocked followed by treatment with vehicle or 100μM dex for a total of four hours. Inset boxes highlight zoomed in regions near the midbody of the animals; white arrowheads denote cytoplasmic and nuclear expression of GFP. Pharyngeal fluorescence is from the co-injection marker *myo-2p::tdTomato;* the scale bar denotes 10 microns. Lower panel bars represent the percentage of mixed-staged, tdTomato-expressing animals with either nuclear only (grey), or cytoplasmic and nuclear expression of GFP after four hours of vehicle or dex treatment (white); * p <0.00001 by a Chi-square test. **(B)** The relative transcript levels of the DAF-16 target gene, *sod-3* after heat shock and ligand treatment for four hours in *daf-2(e1370)* mutants, and wild-type animals carrying a GR LBD::DAF-16A transgene. The graph depicts the mean fold change of expression, relative to vehicle-treated N2 animals lacking any transgenes. Error bars indicate the SEM from at least four biological replicates performed in triplicate (*p<0.03, T-Test).

We next tested whether the GR LBD could similarly modulate the activity of an endogenous *C. elegans* protein. Transcriptionally-inactive DAF-16 is localized to the cytosol, and upon translocation to the nucleus, regulates batteries of genes involved in development, aging, and metabolism (Ogg *et al.* 1997; Lin *et al.* 1997; Henderson and Johnson 2001; Kwon *et al.* 2010; Murphy and Hu 2013). We generated hsp16.48::3XFLAG::GR LBD::DAF-16A transgenic animals, expressed the array through acute heat shock, and assessed the levels of the DAF-16 target gene, *sod-3*, by qRT-PCR. We observed a robust 10.6-fold increase of *sod-3* transcripts after dex treatment, and only a 1.6-fold increase in transcript levels in vehicle-treated transgenics, as compared to wild-type populations (Figure 4B). To compare the levels of *sod-3* upon DAF-16 activation, we also measured *sod-3* levels in *daf-2(e1370)* loss-of-function mutants, which genetically mimic the constitutive nuclear localization and transcriptional activation of DAF-16 (Lin *et al.* 2001; Lee *et al.* 2001). Vehicle and dex-treated populations had 2.8 and 4.3-fold increases in *sod-3* levels, respectively, as compared to wild-type animals. Notably, we only observed significant enrichment of *sod-3* levels upon dex-treatment in *hsp16.48::3XFLAG::GR* LBD::DAF-16A transgenics, and not in *daf-2(-)* mutants or wildtype animals (*p<0.03, T-Test). Together, these results suggest that addition of the GR LBD adduct allows ligand-gated control of DAF-16 activity.

## Conclusions

We found that the QF-GR system was sufficient to drive dex-inducible, tissue-specific expression of a GFP reporter, the toxin gene *peel-1*, and ubiquitous expression of *lin-3c.* We further demonstrate that addition of a GR LBD onto DAF-16 was sufficient to confer ligand-inducible activity of this protein, a potentially generalizable method of regulating the activity and/or localization of nuclear and cytoplasmic proteins in *C. elegans*.

The ability to control gene activation (Logie and Stewart 1995; Metzger *et al.* 1995; Kozlova and Thummel 2002; Banaszynski *et al.* 2006; Cho *et al.* 2013) and protein depletion by small molecule addition (Nishimura *et al.* 2009; Kanke *et al.* 2011; Holland *et al.* 2012; Zhang *et al.* 2015) is a powerful feature in a modern genetic tool box. Our results demonstrate that adding the GR LBD to the QF transcriptional activator confers ligand inducibility to the Q system. While the original Q system could be induced by including the QS repressor and adding quinic acid to relieve repression, it took six hours to detect transgene activity and 24 hours to fully de-repress the system (Wei *et al.* 2012); in contrast, QF-GR allows transgene induction within two hours (Figure 1). Though we were able to observe mRNA induction following dex treatment up to 96-fold over basal expression (Figure 3), we did frequently observe background activity of QF- GR in vehicle treated animals, as discussed below. The QF-GR system should be particularly useful for experiments where heat-shock induction of transgenes is not possible or desirable (i.e., temperature sensitive mutations, physiological effects of heat-shock, etc.), or when sustained transgene expression is required. For example, using QF-GR rather than a heat-shock promoter to drive *lin-3c* would uncouple behavioral lethargy from lin-3c overexpression and lethargy as a result from heat shock (Van Buskirk and Sternberg 2007; Nelson *et al.* 2014).

As a tool, the GR-LBD adduct should be improved and refined going forward. As discussed above, we did frequently observe background activity of QF-GR in vehicle treated animals. Additionally, the GR-LBD did not completely restrict the GR LBD::GFP fusion to the cytoplasm, as would occur in mammalian cells (Picard and Yamamoto 1988). Possible solutions include optimizing position of the GR-LBD relative to a tagged protein, or including two LBD fusions on a construct. Notably, fusing Cre to two ER LBDs in mice conferred tighter regulation and lower background with respect to recombination at floxed alleles (Zhang *et al.* 1996; Casanova *et al.* 2002). We also did not detect transgene expression in the germline. All of our experiments were performed using extrachromosomal arrays, which are frequently silenced in the *C. elegans* germline. However, it is also possible that our failure to observe GFP expression in the germline reflected a failure of dex to enter this tissue. New approaches to license germline expression, such as introns with periodic A/T rich clusters (Frøkjær-Jensen *et al.* 2016) or removal of piRNA binding sites (Zhang *et al.* 2018), combined with singlecopy knock-ins into loci that permit germline expression would distinguish these possibilities. Empirical testing of promoters could also improve the efficacy of GR LBD fusions.

We observed higher basal activity of QF-GR when expressed using the *atf-8* and *eef-1A.1* promoters, as compared to the *egl-17* and *pro-1* promoters (Fig 2B and Sup Fig 1C). In *Drosophila*, progesterone receptor-gated Gal4 was shown to exhibit promoter-specific leakiness in the absence of the RU-486 ligand (Poirier *et al.* 2008), and there is leakiness dependent on the identity of the regulated transgene (Scialo *et al.* 2016). It is unclear what mechanisms drive leaky expression, but some possible explanations are cell-specific differences in proteins that regulate steroid receptors in the presence or absence of ligand, cell- or organism-specific differences in nuclear import/export, or that dex and/or vehicle have unknown QF-GR-independent effects on worm physiology (Picard *et al.* 1990; Freedman and Yamamoto 2004). That being said, all of our experiments utilized extrachromosomal arrays for expression of all transgenes; therefore, we cannot eliminate the possibility that high basal expression from some of the tested constructs was due to high gene dosage from these arrays.

We demonstrated that the GR LBD can be used to confer ligand-gating to three proteins (QF, GFP, DAF-16), highlighting the potential broad utility of this tool in *C. elegans* to regulate protein activity and/or localization. Many intensively studied transcription factors in *C. elegans* (DAF-16, SKN-1, PQM-1, etc.) have their activity regulated by nuclear import/export (Lin *et al.* 2001; Henderson and Johnson 2001; Inoue *et al.* 2005; Tepper *et al.* 2013). Fusing the GR LBD to these factors in transgenes, or knocking the GR LBD into the endogenous locus could confer precise control over protein localization through addition/omission of ligand. The GR LBD would also be useful to add to the recently described cGal toolkit for *C. elegans*, adding an addition level of inducible control (Wang *et al.* 2016). As the cGal system currently lacks the Gal80 repressor, a negative regulator used to repress Gal4 as a “NOT” gate, use of the GR LBD could be a useful alternative NOT gate alleviated by ligand addition (Lee and Luo 1999; Wang *et al.* 2016). The GR LBD could also inactivate cytoplasmic proteins by redirecting them to the nucleus following ligand addition, similar to how the “Anchor-Away” system uses rapamycin-induced dimerization to conditionally inactivate nuclear proteins by shuttling to the cytoplasm (Haruki *et al.* 2008). In theory, with appropriate ligand choice to minimize cross-activation, one could utilize multiple steroid receptor LBDs in one experimental setting to temporally and spatially gate the activity/localization of multiple proteins.

Addition of the GR LBD to regulate protein/activity localization will be a powerful addition to the array of tools to modulate gene expression and protein function in *C. elegans*, including but not limited to CRISPR/Cas9 somatic genome editing (Shen *et al.* 2014), FLP/FRT-mediated gene (in)activation (Voutev and Hubbard 2008), and heterologous gene expression systems (Wei *et al.* 2012; Wang *et al.* 2016).

## Acknowledgements

We thank A. Scacchetti, M. Knuesel, M. Ashina, K. Ehmsen, and the members of the Yamamoto, L’Etoile and Ashrafi labs (UCSF), and the Frand lab (UCLA) for helpful discussions. We also thank C. Van Buskirk (Cal State Northridge), N. Riehs from the Kenyon and Ashrafi labs (UCSF), and X. Wei from the Shen Lab (Stanford) for gifts of plasmids and worm strains. G.C.M was a Robert Black Fellow of the Damon Runyon Cancer Research Foundation (DRG-2189-14) and a UC Presidential Postdoctoral Fellow (PPFP 2013-2015). This work was supported by grants from the National Institutes of Health R01 (CA020535) and R21 (ES026068) to K.R.Y. J.D.W. was supported by the National Institute of General Medical Sciences of the NIH under award K99GM107345 and R00 GM107345. Some strains were provided by the CGC, which is funded by National Institutes of Health Office of Research Infrastructure Programs (P40 OD010440). The content is solely the responsibility of the author and does not necessarily represent the official views of the NIH.

---

**Supplemental Figure 1.**
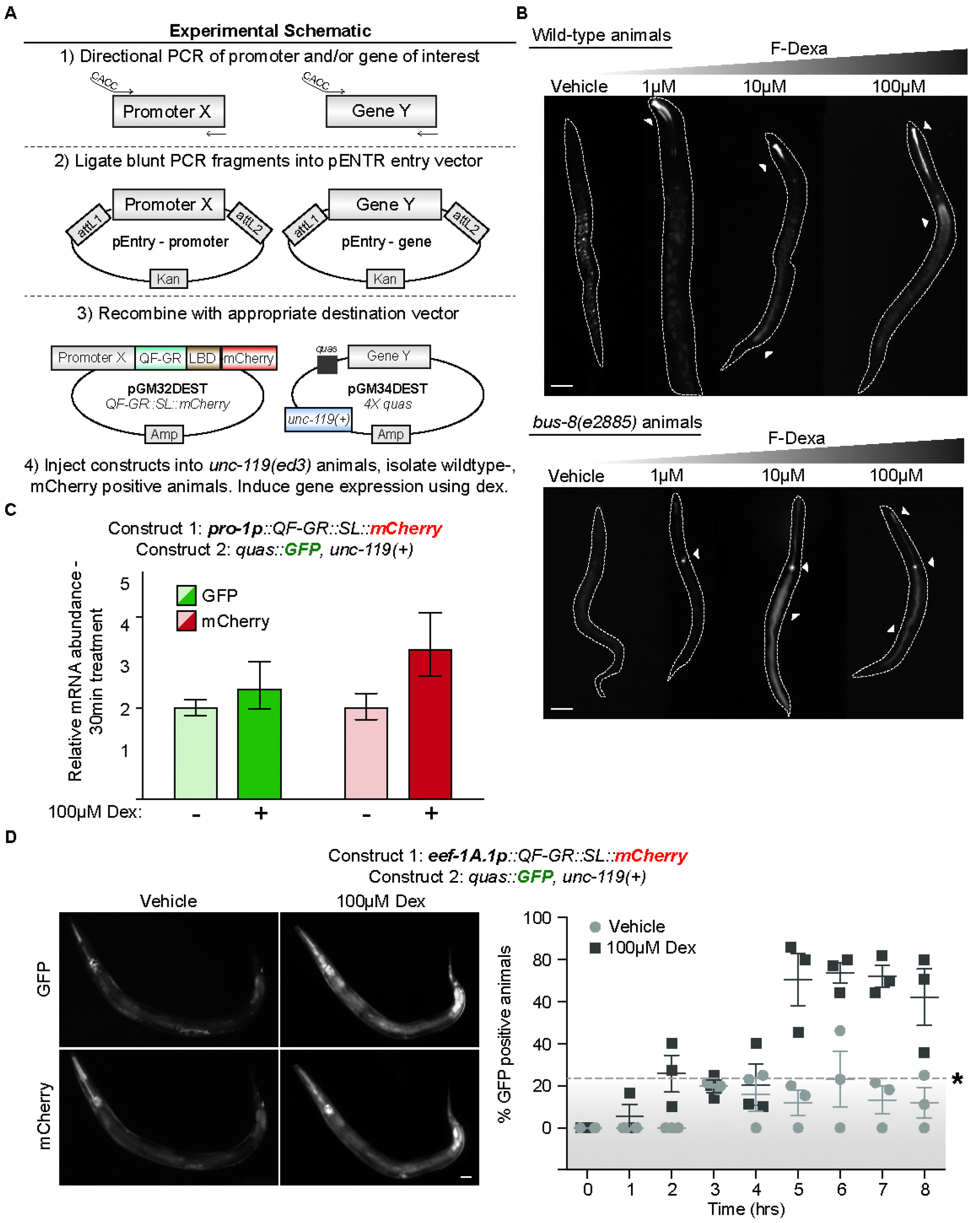
**(A)** Cloning schematic to generate plasmids for gene expression using the dex system. Gateway cassettes upstream of QF-GR coding region, or downstream of the QUAS response element, allow for expression of the gene of interest using the promoter of choice. Using 5’ directional primers, PCR amplifies promoters and target genes from genomic or cDNAs. Blunt PCR fragments are TOPO-cloned into a Gateway entry vector (pEntry, Invitrogen), and the resultant clones are recombined with the appropriate destination vectors, pGM32DEST or pGM34DEST. Both constructs are subsequently injected into *unc-119* mutant animals, and transgenic progeny that are both phenotypically wild-type and red are isolated. Finally, gene induction of the resultant transgenic lines is tested using dex. **(B)** F-Dexa labels the pharyngeal and intestinal lumens of N2 animals and *bus-8* mutants. Representative images of larvae animals incubated with increasing concentrations of F-Dexa or vehicle for two hours. White arrowheads denote lumen fluorescence from F-Dexa; the scale bars denote 20 microns. **(C)** The relative transcript levels of GFP and mCherry after 30 minutes of vehicle or 100μM dex, as measured by qRT-PCR. Bars represent the SEM from three biological replicates performed in triplicate. **(D)** *(Left)* Fluorescent micrographs depict the *eef-1A.1* promoter driving QF-GR expression in representative animals cultivated on either vehicle or 100μM dex plates for 24 hours. Hypodermal and intestinal mCherry expression is observed in both populations. The scale bar represents 20 microns. *(Right)* A time course of GFP-expression of QF-GR driven from the *eef-1A.1* promoter. Mixed-staged animals were cultivated on vehicle or 100μM dex plates, and GFP expression was scored hourly in live animals by fluorescence microscopy. The average percentage of GFP-positive expression is plotted at each time point; error bars represent the SEM of at least three independent trials. N ≥ 31 for each time point; points above the dashed line denote a significant difference from vehicle-treated animals (*p<0.05, T-Test).

